# Proteome trade-off between primary and secondary metabolism shapes acid stress induced bacterial exopolysaccharide production

**DOI:** 10.1101/2024.04.19.590233

**Authors:** Sizhe Qiu, Aidong Yang, Xinyu Yang, Wenlu Li, Hong Zeng, Yanbo Wang

**Author notes:** Corresponding author (H. Zeng); (Y. Wang).

## Abstract

The exopolysaccharide (EPS) produced by *Lactiplantibacillus plantarum* is a high-value bioproduct in food and health industries, and its biosynthesis has been found as a secondary metabolic pathway to mediate acid stress. To quantitatively investigate acid stress response in *L. plantarum* and model EPS production, this study measured metabolomics, proteomics and growth data for *L. plantarum* HMX2 cultured at 4 different pH values. The growth and metabolomics data showed that under acid stress, the EPS production flux was evidently enhanced while the glycolysis and cellular growth were inhibited. The following proteomic analysis found that EPS biosynthetic proteins were significantly up-regulated under acid stress and pinpointed Fur as the most probable transcriptional factor controlling EPS biosynthesis in *L. plantarum*. Furthermore, we identified a proteome trade-off between primary metabolism and EPS biosynthesis, which were then mechanistically depicted by a regulatory proteome constrained flux balance analysis (RPCFBA) model. As the first metabolic model that can simulate secondary metabolism, the RPCFBA model demonstrated good accuracy in predicting growth rates and EPS production fluxes of *L. plantarum* HMX2, validated by experimental data. The *in-silico* perturbation on carbon sources further showed the potential of applying the presented modeling framework to the design and control of microbial secondary metabolism.

## 1. Introduction

*Lactiplantibacillus plantarum* (LP) is a gram-positive lactic acid bacterium found in diverse ecological niches ^1^. LP has been widely used in food and health industries. For instance, it is the major bacterium involved in the fermentation of mozzarella cheese ^2^; it is considered as a safe probiotic that can suppress pathogenic microorganisms ^3^ as well as has immunomodulatory properties ^4^. Many important properties of LP are related to its secreted exopolysaccharide (EPS), which has been extensively studied for its various probiotic effects ^5^, anticancer properties ^6^ and natural ability to improve rheological and sensory properties of fermented foods ^7^. In contrast to primary metabolites, the LP derived EPS is a typical secondary metabolite ^8^, whose biosynthesis directly competes with growth coupled central carbon metabolism for the carbon source and energy. However, the mechanism behind the biosynthesis of EPS has not been well studied compared with its structural and functional properties, thus has long remained murky. The investigation of the relationship between primary metabolism and EPS biosynthesis can provide critical insights and consummate the knowledge of bacterial growth laws ^9^.

In order to achieve precise control of LP derived EPS fermentation, researchers have to elucidate the driving force behind its biosynthesis. Previous studies on the physiological function of lactic acid bacteria derived EPS have indicated that the EPS layer can protect the bacterial cell from acid, osmotic and oxidative stresses ^10^. Furthermore, the significant increase of EPS yield in LP VAL6 strain at low pH found by Nguyen et al. 2021 ^11^ suggests that the EPS biosynthesis is a response to acid stress. In the transcriptomic profiling of LP at pH 5.0, 5.5, and 6.2, some genes involved in EPS biosynthesis (e.g., tagE1, tagE5) were found to be up- regulated for pH 5.0 vs pH 5.5 and pH 6.2 ^12^. However, the up-regulation of EPS biosynthetic genes was not observed for LP CAUH2 strain under oxidative stress ^13^. Subsequently, in our previous work, the analysis of independently modulated gene sets (iModulons) with transcriptomic samples of various conditions (e.g., acid stress, different carbon sources), for the first time, found an interesting trade-off relationship between the regulatory activities of primary metabolism and EPS biosynthesis ^14^. Nevertheless, such trade-off has not been validated by the proteomic profiling of LP under acid stress yet ^15^. In short, various evidence has suggested that the LP derived EPS is a secondary metabolite produced to mediate acid stress, which is beneficial to cell survival as well as renders a long-term population advantage, but the relationship and the regulatory rule between primary metabolism and EPS biosynthesis still remain unclear.

In recent years, computational methods have been increasingly used to assist the design and optimization of the EPS production in lactic acid bacteria. Response surface modeling (RSM), a statistical method, has been widely used to solve the optimal growth medium composition and environmental condition for EPS production ^16,17,18^. Given its broad application, RSM generally treats bacterial metabolism as a “black box”, which makes its results case-specific, therefore limiting its ability for in-depth analysis. Comparatively, flux balance analysis (FBA) with genome-scale metabolic models (GSMMs) is a commonly used “white-box” method to compute metabolic fluxes ^19^. Existing GSMMs of lactic acid bacteria have already included lumped EPS synthetic reactions in their metabolic networks ^20,21^. Nonetheless, several limitations of conventional FBA on secondary metabolism makes it still challenging to model EPS biosynthesis and cellular growth simultaneously ^22^.

To gain an in-depth and quantitative understanding of how acid stress influences primary metabolism and EPS biosynthesis in LP, this study adopted a multi-omics analysis approach to characterize the phenotypic difference of LP HMX2 ^23^ cultured at different pH values, representing different degrees of acid stress, and constructed a regulatory proteome constrained FBA (RPCFBA) model that is capable of simulating cellular growth, central carbon metabolism and acid stress induced EPS biosynthesis simultaneously. More specifically, measured bacterial growth and metabolomics data at 4 different fixed pH values (pH 4.5, 5.0, 5.5 and 6.5) were compared to show how the acid stress affected primary metabolism and EPS production differently. Then, a quantitative proteomic analysis elucidated the acid stress induced regulation of protein expressions by identifying and analyzing significantly up- and down- regulated proteins. Furthermore, the RPCFBA model was constructed and validated with experimental data. Finally, the model demonstrated its practical usefulness in the design and control of microbial secondary metabolism with simulations of growth rates and EPS production under *in-silico* perturbations on carbon sources.

## 2. Results

### 2.1 Inference of EPS structure and characterization of EPS biosynthesis in *Lactiplantibacillus plantarum* HMX2

From the assembled genome sequence of LP HMX2 (NCBI RefSeq assembly, GCF_025144505.1), we predicted the biosynthetic gene cluster of EPS (EPS-BGC), from 1017036 bp to 1063626 bp, that encompassed 20 coding genes using antiSMASH^24^ (**Figure 1A**). The core genes of EPS-BGC were mainly those encoding for glycosyltransferases (GTs) and biosynthetic proteins of activated monosaccharides (e.g., 1_955, UDP-glucose 4- epimerase), which provide precursors for EPS biosynthesis. 1_947, a core gene of EPS-BGC, was found to be a homolog of Wzx flippase for polysaccharide transportation with 67.7% identity. The additional genes contained 3 genes encoding for EPS biosynthetic proteins, that were 1_940/1_953 homologous to cps2/4B and 1_952 homologous to cps2/4A.

**Figure 1.**
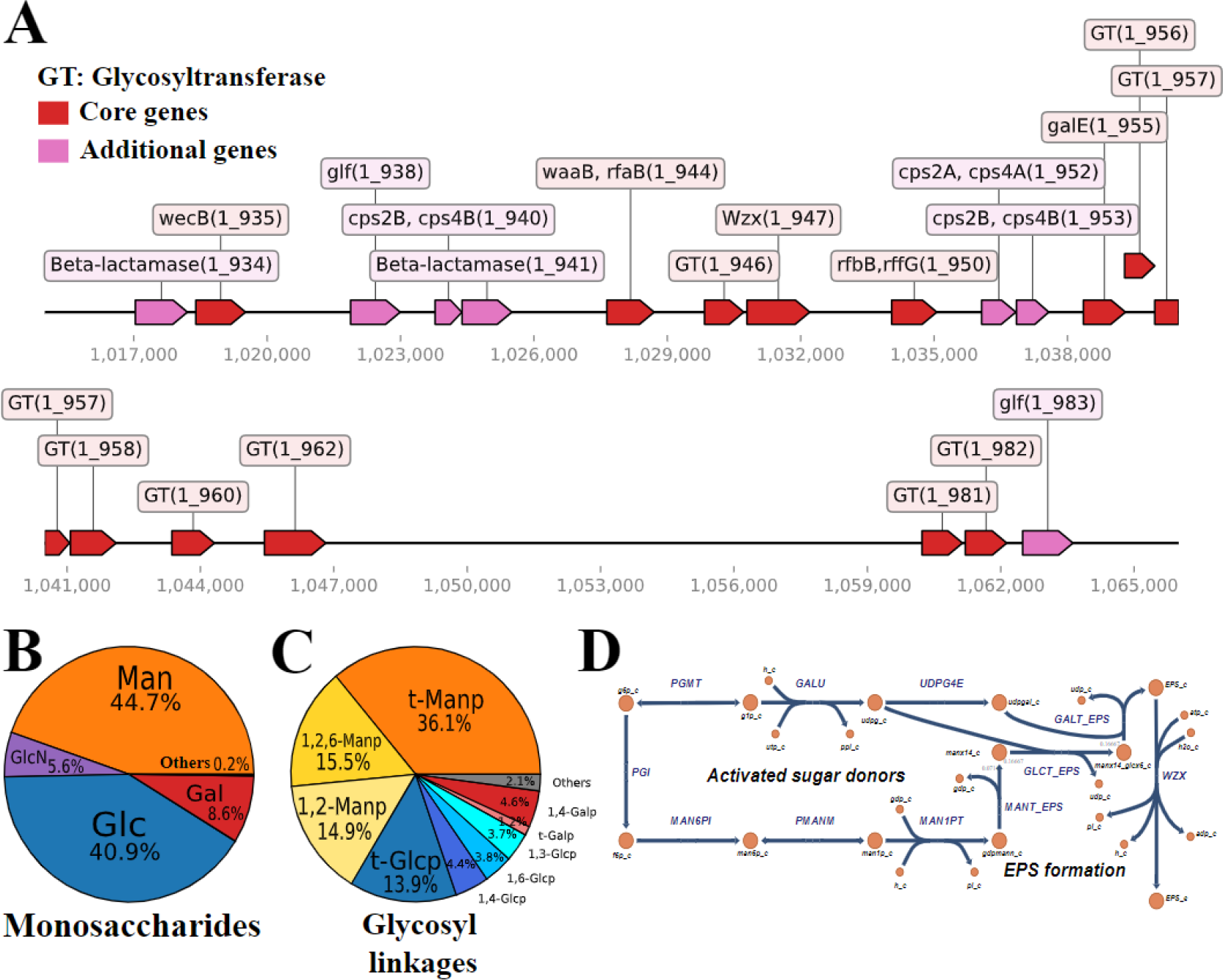
The genome-scale biosynthetic pathway of EPS in LP HMX2. (A) The biosynthetic gene cluster of LP HMX2 EPS. GT: glycosyltransferase, Red: core genes, Pink: additional genes. (B) The molar fractions of monosaccharides in LP HMX2 EPS. (C) The glycosyl linkage compositions of LP HMX2 EPS. (D) The metabolic pathway of EPS biosynthesis. Reaction and enzyme information can be found in **SI, TableS2**. MANT_EPS: mannosyltransferase, GLCT_EPS: glucosyltransferase, GALT_EPS: galactosyltransferase, WZX: Wzx flippase.

The monosaccharide composition of LP HMX2 derived exopolysaccharide (LP-HMX2- EPS) was determined to find out its basic precursors: 44.7% mannose, 40.9% glucose, 8.6% galactose, 5.6% glucosamine and 0.2% other monosaccharides (**Figure 1B**). The high abundance of mannose in LP-HMX2-EPS indicated that the biosynthesis of GDP-mannose using Mannose-6-phosphate isomerase (MAN6PI), Phosphomannomutase (PMANM), and Mannose 1 phosphate guanylyltransferase (MAN1PT) would direct a larger carbon flux from glycolysis than the biosynthesis of other activated monosaccharides. Methylation analysis was conducted to quantitatively determine the glycosyl linkages in LP-HMX2-EPS (**Figure 1C**). The dominance of mannosyl linkages (36.1% t-Manp, 15.5% 1,2,6-Manp and 14.9) suggested that the backbone was mainly composed of 1-2 glycosidic bonds between mannoses while glucosyl and galactosyl linkages likely constituted the branches of the EPS.

Both monosaccharide and glycosyl linkage compositions showed that the EPS mainly consisted of mannose and glucose, but the molar fraction of mannose in glycosyl linkage composition was higher than that in monosaccharide composition. This study considered the glycosyl linkage composition detected by methylation analysis to be more accurate, as the measurement of monosaccharide composition was susceptible to the high glucose concentration in culture media. Subsequently, a pseudo EPS repeating unit was defined (mannose:glucose:galactose = 14:6:1) together with pseudo glycosylation reactions in the GSMM (**Figure 1D**). In this study, we assumed that the composition of EPS was invariant under different conditions.

### 2.2 The influence of acid stress on cellular metabolism

As the pH decreased from 6.5 to 4.5, the growth coupled primary metabolism and EPS biosynthesis in LP HMX2 showed distinctive responses. The LP HMX2 at pH 6.5 reached exponential and stationary phases faster than those at pH 4.5, 5.0 and 5.5 (**Figure 2A**), suggesting that LP HMX2 was a neutrophile. In contrast, the accumulations of EPS at pH 4.5, 5.0 and 5.5 were faster than that at pH 6.5 in the early growth stage (before 4 hr), though the inhibition effects of low pH and undissociated lactic acid ^25,26^ on enzyme activities later hampered the EPS accumulation at pH 4.5 and 5.0 (**Figure 2B**). The computed average growth rates and EPS production fluxes, normalized to unit biomass, revealed that the highest growth rate at pH 6.5 was approximately twice that at pH 4.5 (**Figure 2C**), while the highest EPS production flux at pH 5.0 exceeded twice that at pH 6.5 (**Figure 2D**).

**Figure 2.**
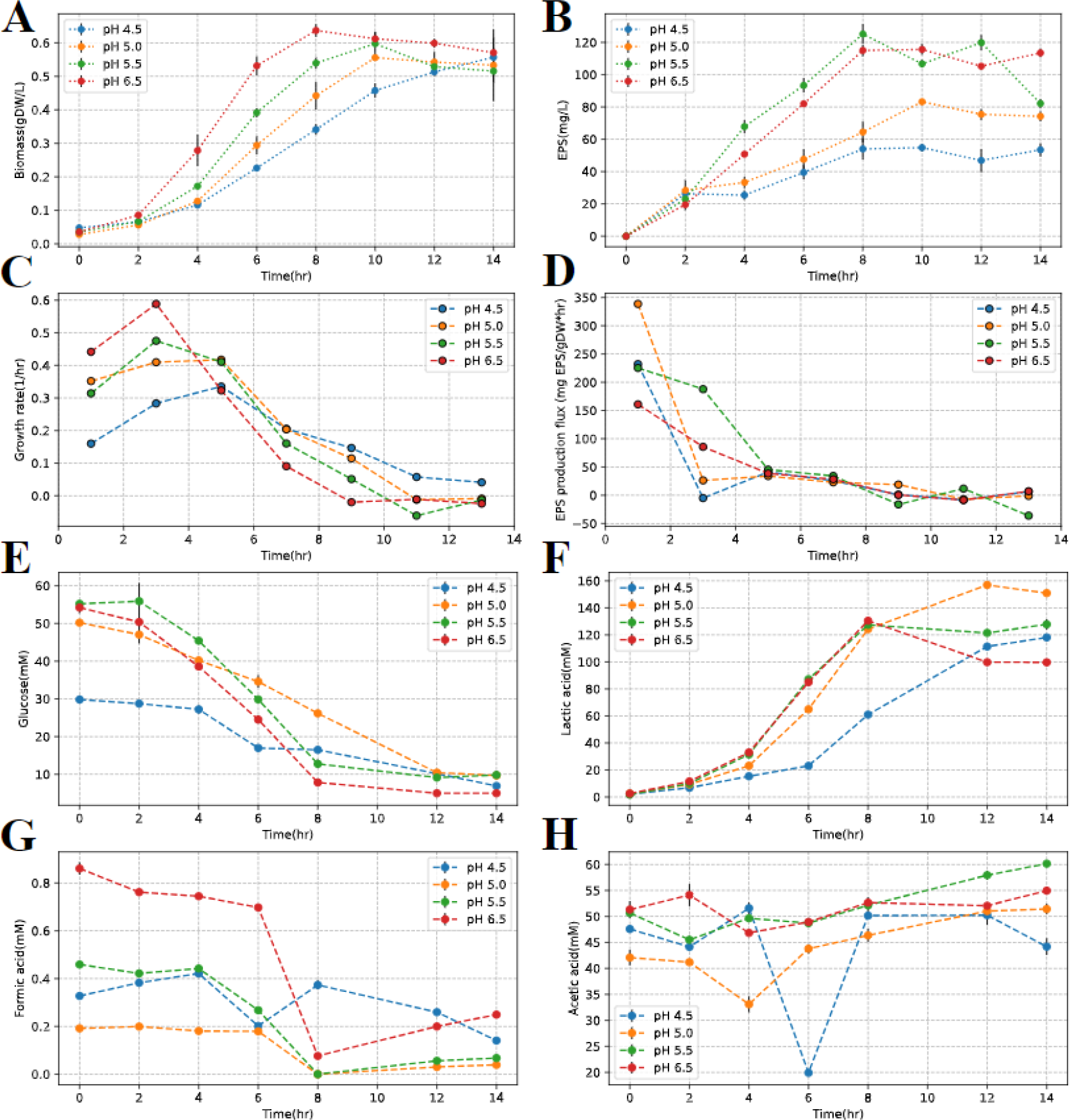
Quantification of primary metabolism and EPS production at fixed pH values of 4.5, 5.0, 5.5, 6.5. (A) The concentrations of cellular biomass converted from OD600. (B) The concentrations of EPS. (C) The average growth rates. (D) The average EPS production fluxes normalized to 1gDW cellular biomass. (E) The extracellular concentrations of glucose. (F) The extracellular concentrations of lactic acid. (G) The extracellular concentrations of formic acid. (H) The extracellular concentrations of acetic acid. The number of replicates was three.

The higher consumption rate of glucose and production rate of lactic acid at pH 6.5 than those under acidic conditions further confirmed that the overall turnover rate of primary metabolism was highest at pH 6.5 (**Figure 2EF**). Apart from indicating the kinetics of primary metabolism, extracellular metabolomics revealed other interesting physiological information of LP HMX2. The decrease of formic acid concentrations in all conditions during fermentation showed that LP HMX2 in MRS media would consume formic acid instead of producing it (**Figure 2G**), possibly for the biosynthesis of purine and pyrimidine ^27^. The consumption rate of formic acid also appeared to be inhibited by the low pH. The relatively stable concentrations of acetic acid in all conditions suggested that LP HMX2 is homofermentative (**Figure 2H**), like most strains of LP ^28^.

The quantification of intracellular metabolomics can provide a direct insight into the metabolic status of LP HMX2 at different pH values. The relatively high concentrations of glycolytic intermediates (i.e., 3-phosphoglyceric acid, phosphoenolpyruvate, pyruvic acid, lactic acid and acetic acid) at pH 5.5, 5.0 and 4.5 indicated the slowdown of central carbon catabolism, caused by the inhibition of glycolytic enzyme activities by the low pH (**Figure 3A-E**). When LP HMX2 entered the stationary phase (12 hr), the high extracellular concentration of undissociated lactic acid (**Figure 2F**) and low pH exhibited a combined inhibitory effect on central carbon catabolism, reflected by the increased intracellular concentrations of glycolytic intermediates. In addition, the relatively high concentrations of formic acid at pH 5.0 and pH 4.5 implied that the biosynthesis of purine and pyrimidine, which consumes formic acid, was inhibited (**Figure 3F**).

**Figure 3.**
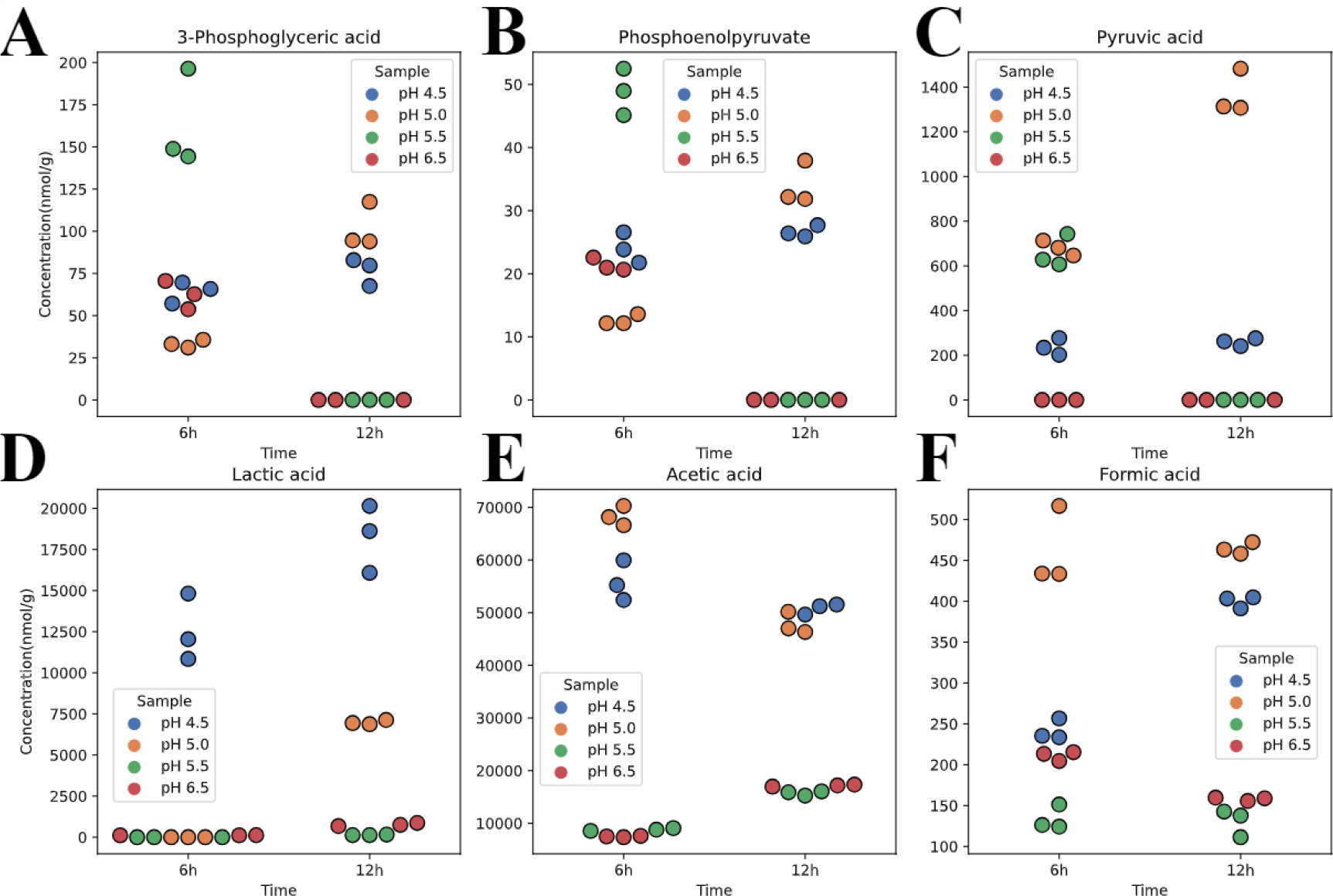
Quantification of intracellular metabolomics at different pH values. (A) The intracellular concentrations of 3-phosphoglyceric acid at 6hr and 12hr. (B) The intracellular concentrations of phosphoenolpyruvate at 6hr and 12hr. (C) The intracellular concentrations of pyruvic acid at 6hr and 12hr. (D) The intracellular concentrations of lactic acid at 6hr and 12hr. (E) The intracellular concentrations of acetic acid at 6hr and 12hr. (F) The intracellular concentrations of formic acid at 6hr and 12hr. The number of replicates was three.

### 2.3 Differential protein expression under acidic conditions

The protein expressions of different pH conditions had observable distinguishing distributions in the principal component analysis (PCA) with high explained variance (PC1: 60.24%, PC2: 17.37%) (**Figure 4A**). The PCA also identified outlier samples at pH 4.5 and pH 5.0, which were replaced with the averages of other samples from the same conditions. With pH 6.5 as the reference condition, differential expression analysis was performed to identify significantly up- and down-regulated proteins (p-value<0.05, absolute log2 fold change(|LFC|)>0.5) (**Figure 4B**). In comparison with the pH 5.5 condition, the amounts of significantly differentially expressed proteins in pH 4.5 and pH 5.0 conditions were much larger. Notably, some up-regulated proteins had extremely large LFCs, whose expressions were presumably induced by acid stress. Regarding proteins encoded by the EPS-BGC (**section 2.1**), significantly up-regulated proteins were only found in pH 4.5 and pH 5.0 conditions (**Figure 4C**), which were mainly GTs (e.g., tagE4 encoded by 1_944, lp_1233 encoded by 1_956) and EPS biosynthetic proteins (e.g., cps4B encoded by 1_953).

**Figure 4.**
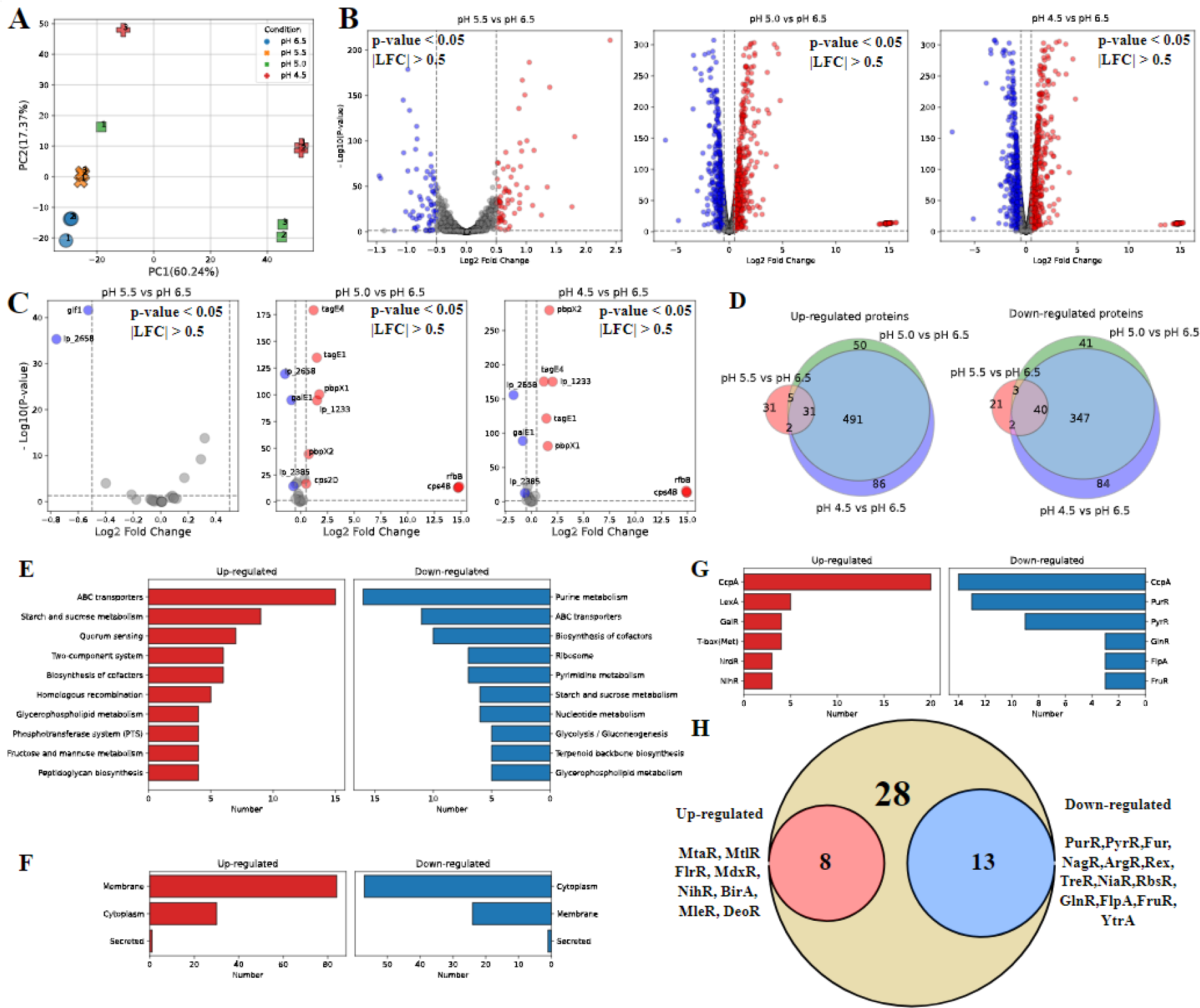
Differential expression analysis of proteomics at different pH values. (A) The PCA biplot of proteomics data. (B) Volcano plots of differential expression analysis of pH 5.5 vs pH 6.5, pH 5.0 vs pH 6.5 and pH 4.5 vs pH 6.5. LFC: log2-fold change. (C) Volcano plots of differential expression analysis for proteins encoded by the EPS BGC of pH 5.5 vs pH 6.5, pH 5.0 vs pH 6.5 and pH 4.5 vs pH 6.5. (D) Venn diagrams of up- and down-regulated proteins of pH 5.5 vs pH 6.5, pH 5.0 vs pH 6.5 and pH 4.5 vs pH 6.5 (p-value<0.05, |LFC|>0.5). (E) KEGG pathway enrichment analysis of differentially expressed proteins (p-value<0.05, |LFC|>0.5). (F) Subcellular location enrichment analysis of differentially expressed proteins (p-value<0.05, |LFC|>0.5). (G) Regulon enrichment analysis of differentially expressed proteins (p-value<0.05, |LFC|>0.5). (H) Up- and down-regulated transcription factor proteins under acidic conditions (p-value<0.05, |LFC|>0.5). The p-value was adjusted with Benjamini-Hochberg method. The number of replicates was three.

Samples at pH 4.5 and 5.0 were considered as under acid stress, because the significantly differentially expressed proteins in those two conditions had much higher consistency with each other than with those at pH 5.5 (**Figure 4D**). The proteins that were significantly up- or down-regulated at both pH 4.5 and 5.0 were extracted to perform the enrichment analysis for functional pathways, subcellular locations, and regulons. The up- regulated proteins enriched in ATP-binding cassette (ABC) transporters, carbohydrate metabolism, and stress response, while the down-regulated proteins enriched in purine metabolism, ABC transporters, and biosynthesis of cofactors (**Figure 4E**), which were highly consistent with previous research on acid stress induced gene expression regulation of LP^12,14,15^. For subcellular locations, most up-regulated proteins (e.g., ABC transporters, stress response proteins, EPS biosynthetic proteins) were membrane proteins, while most down- regulated proteins (e.g., enzymes in purine and pyrimidine metabolism) were located in the cytoplasm (**Figure 4F**).

With regulons of LP obtained from RegPrecise^29^, both up- and down-regulated proteins were found to be enriched in CcpA regulon (**Figure 4G**). Five stress response proteins encoded by genes in LexA regulon were significantly up-regulated. Most proteins encoded by genes in PyrR and PurR regulons were significantly down-regulated. Among 49 identified transcriptional factors (TFs) of LP, 8 and 13 TFs were significantly up- and down-regulated under acid stress, respectively (**Figure 4H**). The significant down-regulation of PyrR and PurR was in agreement with the enrichment analysis of functional pathways and regulons. For TFs controlling nutrient uptake, significantly down-regulated GlnR and FruR are repressors of glutamine ABC transporter and fructose-specific PTS, respectively, while up-regulated MtlR and MdxR are activators of mannitol-specific PTS and maltose/maltodextrin ABC transporter, respectively.

Regarding the regulation of EPS biosynthesis, Fur was found to be significantly down-regulated under acid stress, in agreement with previous studies suggesting that Fur directly or indirectly represses EPS biosynthesis ^30,31,32^ and mediates acid tolerance ^33,34^. Besides, the systematic transcriptomic analysis of LP also indicated that Fur had significant correlations with acid stress and EPS biosynthetic genes ^14^.

### 2.4 Acid stress induced proteome resource allocation

With gene-reaction-protein rules in the GSMM of LP HMX2, the protein expression level changes of metabolic reactions between pH 4.5 and pH 6.5 were computed to investigate how the proteome resource allocation of metabolic pathways was affected by acid stress (**Figure 5**). At pH 4.5, most glycolytic enzymes were only slightly down-regulated compared to pH 6.5, except that L-lactate dehydrogenase (LDH_L) had a LFC of -0.78. Different from other glycolytic enzymes, Glyceraldehyde-3-phosphate dehydrogenase (GAPD) was significantly up-regulated. In the pentose phosphate pathway (PPP), Phosphogluconate dehydrogenase (GND) and Ribulose 5-phosphate 3-epimerase (RPE) were significantly down-regulated, while Transaldolase (TALA) was significantly up-regulated. For purine and pyrimidine biosynthesis, most enzymes were significantly down-regulated, particularly Dihydroorotase (DHORTS), Dihydoorotic acid dehydrogenase (DHORD) and Orotate phosphoribosyltransferase (ORPT). In general, acid stress resulted in the reduction of proteome resources allocated to glycolysis and the biosynthesis of DNA/RNA materials.

**Figure 5.**
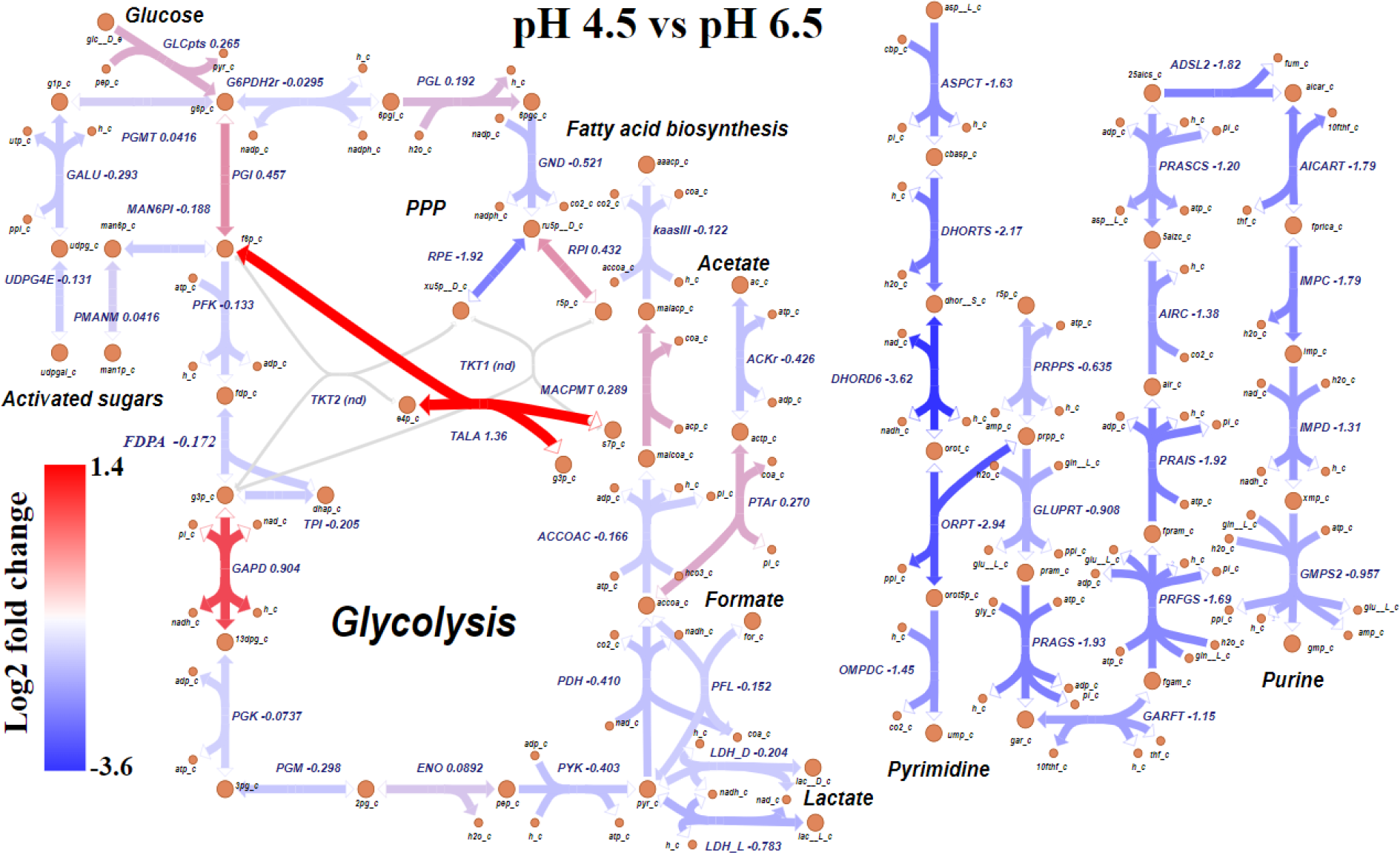
Protein expression level changes (pH 4.5 vs pH 6.5) of metabolic reactions in central carbon metabolism and pyrimidine/purine metabolism. Detailed reaction and enzyme information can be found in **SI, TableS2**.

In the theoretical model of proteome resource allocation proposed in this study, the proteome of LP HMX2 was divided into an inflexible housekeeping sector (Q) and four flexible sectors (A, C, T, U) (**section 4.6**). The C sector included all glycolytic enzymes; the T sector included glucose uptake via phosphoenolpyruvate-dependent sugar PTS and acid transporters; the A sector included all ribosomal and chaperone proteins; and the U sector contained all proteins encoded by the EPS-BGC (**Figure 6A**). The mass fractions of four flexible sectors at different pH values were computed with protein expression levels (**Figure 6B**). The fraction of the C sector dropped substantially when the pH decreased from 5.5 to 5.0, but the declining trend then became smaller from 5.0 to 4.5. The fraction of the A sector dropped when the pH decreased from 6.5 to 5.5, but only had slight decreases for lower pH values. The fraction of the T sector increased continuously with the decrease of pH, which indicated that the inhibition of transporter activity at low pH demanded the cell to reallocate more proteome resources for maintaining the uptake of nutrients. When the pH decreased from 5.5 to 5.0, the fraction of the U sector had a ∼2-fold increase, in agreement with the observed ∼2-fold increase of the EPS production flux when the pH decreased from 6.5 to 5.0 (**Figure 2D**).

**Figure 6.**
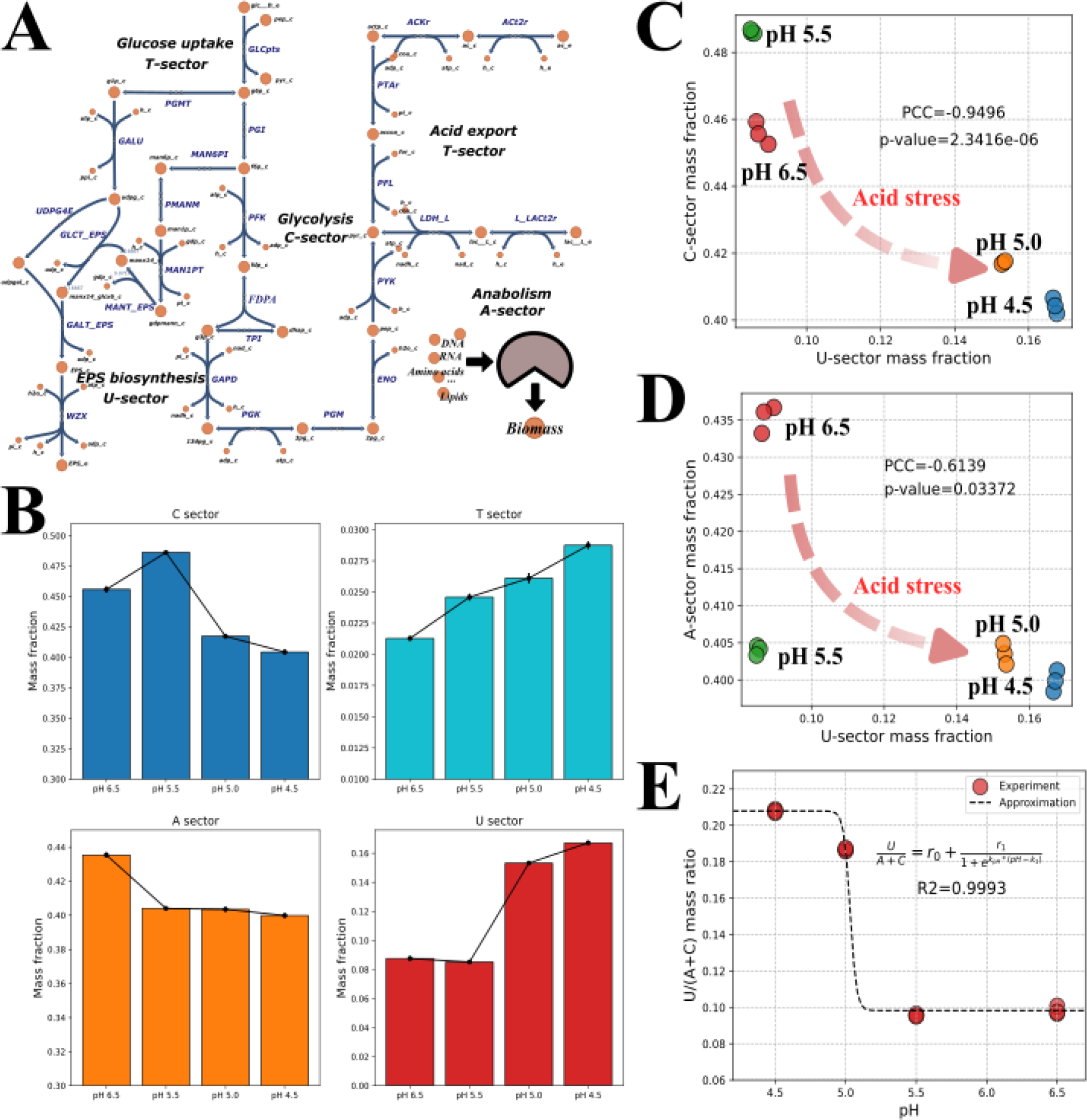
Proteome resource allocation among functional sectors under acid stress. (A) Illustration of flexible proteome sectors in the proteome resource allocation model. A: anabolism; T: transportation; C: catabolism; U: EPS biosynthesis. (B) Proteome sector mass fractions at different pH values. (C) The Pearson correlation of C and U sectors (p-value<0.05). (D) The Pearson correlation of A and U sectors (p-value<0.05). (E) The numerical approximation of as a sigmoid function of pH values (R2=0.9993). =0.0876, =0.07953, =44.9457, =5.0344.

Furthermore, the significantly negative correlations of A and C sectors with the U sector depicted an explicit proteome trade-off between primary and secondary metabolism (**Figure 6CD**), which was also reported in the elucidation of independently modulated genes (iModulons) in LP ^14^. When the pH decreased from 6.5 to 4.5, the proteome resource occupied by energy metabolism and anabolism decreased, while more proteome resources were allocated to EPS biosynthetic pathway. In addition, a sigmoid function was employed to approximate the variation in the mass fraction of the U sector in response to the pH decrease (**Figure 6E**), and was subsequently incorporated into the proteome constrained FBA to simulate the EPS production flux (**section 4.6**, **Eq. 7**).

### 2.5 Simulation of primary metabolism and EPS production

Based on measured metabolomics data (**section 2.2**) and acid stress induced proteome resource allocation observed in this study (**section 2.4**), a regulatory proteome constrained FBA (RPCFBA) model was constructed to simulate the primary metabolism and the EPS production flux of LP HMX2. The simulation predicted the maximum growth rates, EPS production and glucose uptake fluxes at different pH values ranging from 4.5 to 6.5, without considering the inhibition of undissociated lactic acid. The simulation results were compared with the experimental data to evaluate the accuracy of model predictions.

When pH decreased from 6.5 to 5.5, the simulated growth rate consecutively dropped from 0.79 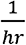 to 0.61 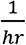 due to the decrease of enzyme activities and the increase of 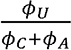 (**Figure 7A**). Oppositely, the EPS production flux was enhanced by the increase of the proteome resource allocated to the EPS biosynthetic pathway (**Figure 7B**). Around pH 5.0, the sudden jump in the sigmoid function of 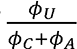 (**Figure 6E**) led to the sudden decrease of simulated growth rate and sudden increase of EPS production flux (**Figure 7AB**). When the pH continued to decrease below 5.0, the simulated EPS production flux decreased, as the activities of EPS biosynthetic enzymes decreased, but the simulation predicted that the growth rate would slightly increase because the carbon flux and energy (activated sugars) consumed by EPS biosynthesis were reduced. However, the experimental growth rate at pH 4.5 did not become higher than that at pH 5.0, which could be resulted from the inaccuracy of the pH-dependency of enzyme activities parameterized in this model. The trend of simulated glucose uptake fluxes was close to that of simulated growth rates, but there appeared to be an observable inconsistency between simulated and experimental glucose uptake rates (**Figure 7C**). Such inconsistency possibly resulted from the underrating of the energy demand (ATP) for growth in the GSMM. In general, the simulated growth rates and EPS production fluxes were close to the experimental data (R2=0.579 for growth rates, R2=0.775 for EPS production fluxes).

**Figure 7.**
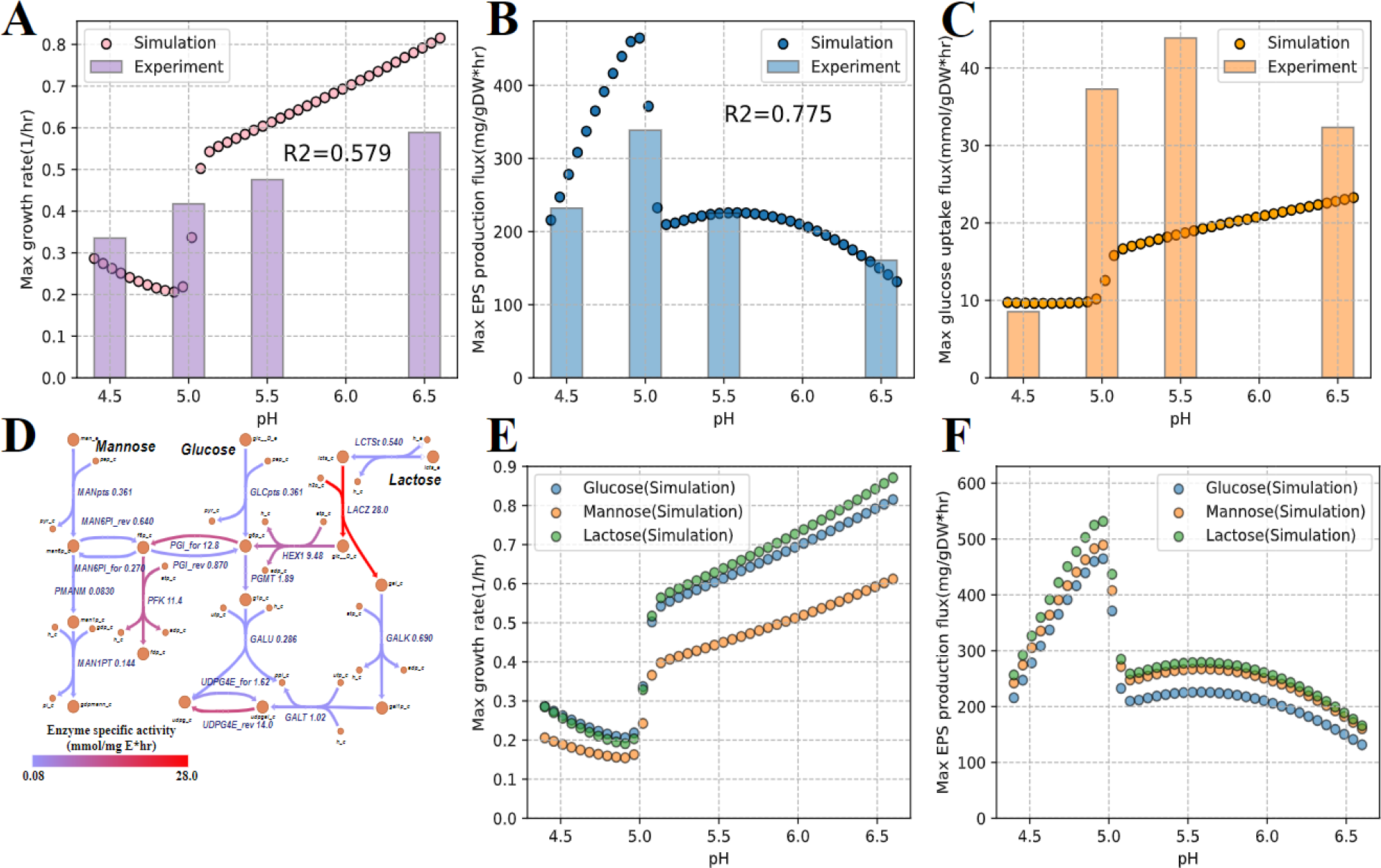
The simulation of primary metabolism and EPS production at different pH values. (A) The simulated and experimental growth rates (—). (B) The simulated and experimental EPS production fluxes (—). (C) The simulated and experimental glucose uptake fluxes (—). (D) Enzyme activities of metabolic pathways for glucose, lactose and mannose. The details of enzyme activities can be found in **SI, Table S3**. (E) The simulated growth rates with different carbon sources. (F) The simulated growth rates with different carbon sources.

Previous studies have indicated that different carbon sources can influence the EPS production rate of lactic acid bacteria ^35,36^. Therefore, we conducted an *in-silico* perturbation on the carbon source of the growth medium, where the sole carbon source was set as glucose, mannose or lactose, respectively. Similar to glucose, mannose can also be uptake via PTS, as the proteome of LP HMX2 contained PTS subunits specific for mannose (1_479, 1_480, and 1_481), but the low activity of Mannose-6-phosphate isomerase (MAN6PI) (in contrast to Glucose-6-phosphate isomerase (PGI)) will theoretically increase the proteome cost of glycolysis (**Figure 7D**). For lactose, LP needs to first break it down to glucose and galactose, and thus requires two more steps than utilizing glucose (**Figure 7D**). However, the activities of Hexokinase (HEX) and Beta-galactosidase (LACZ) are high. With respect to EPS biosynthesis, the change of carbon source from glucose to lactose or mannose can directly provide galactose or mannose, which will reduce the proteome cost of using Mannose-6-phosphate isomerase (MAN6PI), Phosphoglucomutase (PGMT), UTP-glucose-1-phosphate uridylyltransferase (GALU) and UDPglucose 4-epimerase (UDPG4E) to synthesize UDP-galactose and GDP-mannose from glucose. As expected, the simulation predicted that the maximum growth rates of using glucose and lactose were close, while the maximum growth rates of using mannose were noticeably lower (**Figure 7E**). In contrast, the EPS production fluxes of using mannose and lactose were predicted to be higher than that of using glucose (**Figure 7F**). In conclusion, our *in-silico* perturbation study predicted that lactose will be the most favorable carbon source, compared to glucose and mannose, for optimal bacterial growth and EPS productivity.

## 3. Discussion

Although the high value of LP derived EPS has been demonstrated in various areas, the mechanistic understanding of the driving force behind its biosynthesis still remains to be uncovered. The observation that EPS production can be enhanced by a low pH in a recent study ^11^, together with the development of proteome allocation theory ^22,37,38^, shed light on resolving this issue. This study confirmed that the EPS production flux indeed increased under acidic conditions. Subsequently, the proteomic analysis found that EPS biosynthetic proteins were significantly up-regulated by acid stress. It also identified the most probable TF, namely Fur, regulating the expression of EPS biosynthetic proteins (**section 2.3**). Unfortunately, due to the lack of knowledge on transcription units of LP, this study currently cannot derive a detailed regulatory mechanism for proteins encoded by the EPS-BGC. Nevertheless, the analysis of expression levels of functional proteome sectors quantitatively identified an acid stress induced proteome trade-off between primary metabolism and EPS biosynthesis in LP HMX2 (**section 2.4**), which unveiled the driving force behind EPS production in lactic acid bacteria.

To quantitatively describe the trade-off between primary and secondary metabolism, existing theoretical models such as Grime’s competitor-stress-ruderal triangle ^39,40^ and Synthetic Chemostat Model ^38^ have made great contributions to propose a resource allocation framework, but none of them has been implemented to model metabolic fluxes yet. The RPCFBA model, constructed based on multi-omics data in this study, formulated the acid stress induced proteome resource allocation into a concise mathematical function (Eq. 6), and accurately predicted the changes of growth rates and EPS production fluxes in response to the decrease of pH (**section 2.5**). Also, the *in-silico* perturbation study on carbon sources demonstrated the broad potential and practical usefulness of RPCFBA on designing and optimizing complex fermentation processes, especially those involving secondary metabolisms for producing high-value bioproducts.

Nonetheless, some limitations remain in the presented modeling framework. The large deviation between experimental and simulated glucose uptake rates (**Figure 7C**) indicated that the energy demand (ATP) for growth in the GSMM might need to be re-evaluated for the specific LP strain, though the inaccuracy in the measurement of glucose concentrations was also a possible error source. Another weakness of the current model is its inability to accurately represent pH dependent changes in enzyme activities. Enzyme *k*_Cat_ predictors considering pH might resolve this issue, but the existing most accurate predictor can only achieve a RMSE of 0.594 ^41^. Most importantly, the RPCFBA model cannot yet account for the inhibition of undissociated lactic acid on cellular growth of LP, thus is incapable of simulating the decrease of growth rate in time when the carbon source is abundant. The experimental data of this study reflected that the empirical function used in Özcan et al., 2021 ^42^ and Qiu et al., 2023 ^43^ to model the inhibition of undissociated lactic acid is not universally applicable. The growth rates at constant pH 6.5 and pH 5.5 dropped fast in time (**Figure 2C**), though the theoretical concentrations of undissociated lactic acid should be small based on Henderson-Hasselbalch equation. Consequently, a better method to model the complicated inhibitory effect of undissociated lactic acid on growth needs to be developed to enhance the performance of metabolic modeling of lactic acid bacteria grown in complex and changing conditions.

In a nutshell, this study provides insights into the growth strategy of LP HMX2 under acid stress, particularly on how it balances growth and stress response via reallocating its proteome resources between primary and secondary metabolism. In future studies, determining transcription units on the genome of LP and conducting ChIP-seq experiments for TFs associated with acid stress response (e.g., Fur) might uncover a detailed regulatory signaling pathway of how acid stress activates the expression of EPS biosynthetic proteins. The RPCFBA model established in this study can satisfactorily capture the changes of growth rates and EPS production fluxes induced by acid stress, though the quantitative accuracy can be further improved. With further refinement and tailored modifications, we can envisage that the RPCFBA model can potentially become a generic modeling framework for the design and control of biosynthesis of various valuable secondary metabolites associated with stress response (e.g., antibiotics produced by actinomycetes ^44^).

## 4. Materials and Methods

### 4.1 Strains, media, and culture conditions

*Lactiplantibacillus plantarum* HMX2 (LP HMX2), an EPS producing strain isolated from Chinese Northeast Sauerkraut ^23^, was used to investigate the effect of acid stress on the secondary metabolism of lactic acid bacteria. For activation, 3% inoculum of LP HMX2 was introduced into De Man, Rogosa and Sharpe (MRS) broth and incubated anaerobically at 37°C for 12 hours in a sterile laminar flow hood. For culture experiments, 10ml activated bacterial culture was inoculated into a conical flask with 3L MRS broth, sealed with a membrane. The temperature was maintained at 37°C. The LP HMX2 was cultured at 4 different fixed pH values: 6.5, 5.5, 5.0, and 4.5, and each condition had 3 replicates. The pH of each condition was adjusted to and maintained at the fixed value using titration with sodium hydroxide and sulfuric acid, tracked by the fermentation monitor (iCinac, AMS, France) with an Inlab Smart pro-ISM detection electrode (Mettler Toledo, Switzerland) (**see SI, Figure S1**). The pH 6.5 was selected as the reference condition in metabolomic and proteomic analysis, as it is the upper bound of optimal growth pH for most LP strains ^45^. The culture experiment lasted for 14 hours for each condition and samples of 100ml liquid culture media were taken every 2 hours for further analysis. The OD600 was measured using a spectrophotometer at 600 nm for the quantification of growth kinetics, and converted to dry weight biomass concentrations with a conversion factor of 0.35 ^gDW^ OD_600_ ^46^. The concentration of produced EPS was estimated by the phenol-sulfuric acid method ^47^ using glucose as standard, and subtracting the amount determined at zero time. The measurement of metabolites (e.g., glucose, lactic acid, etc.) was elaborated in **section 4.3**.

### 4.2 Whole genome sequencing analysis

The genome sequencing of LP HMX2 was performed using the Hiseq/Novaseq sequencing platform. The genomic DNA was extracted using the DNeasy blood and tissue kit (Qiagen, Hilden, Germany), and then fragmented into a size within 500bp using Bioruptor Sonication Device (Covaris S220). End Prep Enzyme Mix was used for the end repair of fragmented DNA sequences. A HiFi sequencing library for de novo genome assembly was prepared using the SMRTbell library prep kit (PacBio, Menlo Park, CA, USA), according to the manufacturer’s protocol. Then, the sequencing was carried out using Illumina HiSeq instrument (Illumina, San Diego, CA, USA) following ABySS v. 12.1 assembly method. After reads with lengths smaller than 500bp were filtered out, HGAP4/Falcon ^48^ was used to assemble the cleaned reads.

The coding genes on the assembled genome of LP HMX2 were identified using Prodigal ^49^, and annotated with various databases including Non-Redundant Protein Database (NR) ^50^, Kyoto Encyclopedia of Genes and Genomes (KEGG) ^51^, Clusters of Orthologous Genes (COG) ^52^, Carbohydrate-Active enZYmes (CAZy) ^53^ and Transporter Classification Database (TCDB) ^54^.

The locus tags of coding genes in LP HMX2 can be found in https://github.com/SizheQiu/LbPtEPS/tree/main/data/Genome_HMX2. The biosynthetic gene cluster (BGC) of EPS was predicted using antiSMASH bacterial version ^24^.

### 4.3 Quantification of intra- and extra-cellular metabolomics

The concentrations of intra- and extra-cellular metabolites were measured to quantify metabolic exchange fluxes (uptake and secretion) and intracellular metabolic status. For extracellular metabolites, samples were filtered through a 0.22 μm membrane and the supernatant was collected. For intracellular metabolites, samples were centrifuged at 9000 rpm for 10 minutes. Then, the pellet was collected and washed with the PBS buffer. The extracellular metabolomic data can be found in https://github.com/SizheQiu/LbPtEPS/blob/main/data/Exp_data/Metabolomics_mM.csv, and the intracellular metabolic data can be found in https://github.com/SizheQiu/LbPtEPS/blob/main/data/Exp_data/IntraMetabolomics.csv.

The concentrations of amino acids, glucose, lactic acid and glycolytic intermediates (except glucose-6 phosphate and fructose 6-phosphate) were measured using LC/MS: UPLC (LC-30AD) equipped with a MS-8050 triple quadrupole mass spectrometer. To prepare the samples, 600 μL of acetonitrile was added to 200 μL of test and QC samples separately. Then, each sample was vortexed and centrifuged with 12000 RCF at 4L for 10 min, and the supernatant of each sample was diluted at 4-fold, 20-fold and 100-fold with water. 20 μL of internal standard solution was added to each sample. Next, each sample was vortexed with 1000 RCF at 25 L for 10 min, and centrifuged with 4680 RCF at 4L for 10 min. The columns used in LC were ACQUITY UPLC HSS T3 and Discovery HS F5 HPLC, and the temperature was 45L. The injection volume was 1uL, and the flow rate of the mobile phase (A=0.1% formic acid/water; B=0.1% formic acid/acetonitrile) was 0.3 mL/min. The gradient elution procedure was as follows: 0 min, 100% A; 3 min, 100% A; 7 min, 40% A, 60% B; 8 min, 5% A, 95% B; 10 min, 5% A, 95% B; 10.1 min, 100% A. The acquisition mode of MS was MRM. The interface voltage was 4.0 kV. The temperatures of the interface, desolvation line and heater block were 300 L, 250 L and 400 L, respectively. The flow rates of automatic gas, drying gas and heating gas were 3.0 L/min, 10.0 L/min and 10.0 L/min.

For glucose-6 phosphate and fructose 6-phosphate, the sample preparation and the LC/MS equipment were the same. The column of LC was ACQUITY UPLC BEH Amide with a temperature at 45L. The injection volume was 1uL, and the flow rate of the mobile phase (A= 20 mM ammonium acetate, 1.2% ammonia, 95% water, 5% acetonitrile; B= acetonitrile) was 0.3 mL/min. The gradient elution procedure was as follows: 0 min, 25% A, 75% B; 1 min, 40% A, 60% B; 10 min, 40% A, 60% B; 13 min, 25% A, 75% B; 13.1 min, 25% A, 75% B. The interface voltage in MS was set at 4.0 kV, and the other settings in MS were the same as those in LC/MS for other metabolites, as described above.

The concentrations of short-chain fatty acids and acetic acids were measured using GC/MS: GC-2030 equipped with a TQ 8040NX mass spectrometer. For the sample preparation of fatty acids, 20 μL of 5% aqueous phosphoric acid solution was added to 50 μL of the sample, and each sample was vortexed for 5 min. Then, 100 μL of methyl tert-butyl ether was added, and each sample was vortexed for 5 min and centrifuged with 12000 rpm/min for 5 min at 4 L. The 70 μL of supernatant was transferred to the injection vial. For the sample preparation of acetic acid, 20 μL of 5% aqueous phosphoric acid solution was added to 50 μL of the sample, and each sample was vortexed for 5 min. Then, 100 μL of methyl tert-butyl ether was added, and each sample was vortexed for 5 min and centrifuged with 12000 rpm/min for 5 min at 4 L. The 5 μL of the supernatant was added into 495 μL of methyl tert-butyl ether, and transferred to the injection vial. For the settings in GC, the column was DB-FFAP (30 m*0.32 mm*0.25 μm), the injection volume was 1 μL, the flow rate was 1 mL/min, the loading gas was helium, and the temperature was 230 L. The gradient elution procedure was as follows: 0 L/min, 60 L, 2 min holding time; 2 L/min, 100 L, 0 min holding time; 5 L/min, 110 L, 0 min holding time; 10 L/min, 160 L, 0 min holding time; 20 L/min, 240 L, 2 min holding time. For the settings in MS, the acquisition mode was Q3 SIM, the source temperature was 230 L, the interface temperature was 200 L, and the detector voltage was 1.04 kV.

### 4.4 Structural analysis of LP-HMX2 derived exopolysaccharide

The isolation and purification of LP HMX2 derived exopolysaccharide (LP-HMX2-EPS) followed the experimental procedure in Yang et al., 2018 ^55^. Then, the monosaccharide and glycosyl linkage compositions of the purified LP-HMX2-EPS were quantitatively determined (**see SI, Table S1**), and were used to build pseudo-reactions of EPS biosynthesis in the modified GSMM (**see SI, Supplementary method 1.1**).

The mass fraction of each monosaccharide in LP-HMX2-EPS was measured using LC/MS: ThermoU3000 HPLC system. The 5 mg purified EPS sample was added to 1 ml 2 M trifluoroacetic (TFA) acid solution, and heated at 121 °C for 2 hours. The sample was washed with 3 ml methanol and dried with nitrogen gas for 3 times. The dried sample was dissolved in 5ml sterile deionized water. 0.2 ml 0.5 M NaOH and 0.5 ml 0.5 M PMP-methanol were added to 0.2 ml sample to react at 70 °C for 1 hour to form sugar-PMP derivatives. 0.2 ml 0.5 M HCl was added to neutralize NaOH. Vortex-assisted extraction with chloroform was used to remove excess PMP. The sample volume was adjusted to 1 ml with sterile deionized water before detection. For the settings in ThermoU3000 HPLC system, the column was ZORBAX EclipseXDB-C18, the flow rate of the mobile phase (acetonitrile:12 g/L monobasic potassium phosphate/2 M NaOH=17:83) was 0.8 mL/min for isocratic elution, the column temperature was 30 °C, the detection wavelength was 250 nm, and the injection volume was 10 μl. The mass fractions of mannose, glucosamine, rhamnose, glucuronic acid, galacturonic acid, galactosamine, glucose, galactose, xylose, arabinose and fucose were quantified using external standard calibration.

Methylation analysis and GC/MS (Agilent 7890A GC system equipped with Agilent 5977B quadrupole mass spectrometer) were used to determine the glycosyl linkage composition through the quantification of different monosaccharide residues (e.g., 1,2-Manp). The 10 mg EPS sample was dissolved in 1 ml deionized water, and 1 ml 100 mg/ml carbodiimide was added to react for 2 hours. 1ml 2M imidazole and 1 ml 30 mg/ml sodium borodeuteride (NaBD4) were added to react for 3 hours. 100 μL acetic acid was added to terminate the reaction. The sample was filtered and freeze-dried. Then, the methylation of LP- HMX2-EPS followed the experimental procedure in Ciucanu & Kerek, 1984 ^56^. Next, the methylated sample was injected into the Agilent 7890A-5977B GC/MS system. The column was HP-5MS capillary column (30 m × 0.25 mm × 0.25 μm, Agilent J&W Scientific, Folsom, CA, USA), the loading gas was helium, the flow rate was 1.0 mL/min, the injection volume was 1μL, and the temperature of the injection port was 260 °C. The gradient elution procedure was as follows: 0 L/min, 50 L, 1 min holding time; 50 L/min, 130 L, 3 L/min, 230 L, 2 min holding time. The Agilent 5977B quadrupole mass spectrometer was equipped with an electron impact ion source and a MassHunter workstation. The temperature of the injection port and the quadrupole were 230 L and 150 L respectively. The electron energy was 70 eV. The scan mode was full scan with a range of m/z 30 to m/z 600.

### 4.5 Quantitative proteomic analysis

For each condition, samples were taken at the exponential phase (i.e., 6 hr), and SDT (4% SDS, 100 mM Tris-HCl, pH 7.6) buffer was used for sample lysis and protein extraction. 20 µg of protein for each sample was mixed with the 5X loading buffer and boiled for 5 min, and then separated on 4%-20% SDS-PAGE gel (constant voltage 180 V, 45 min). Protein digestion by trypsin was performed following filter-aided sample preparation (FASP) procedure ^57^. The digested peptides of each sample were desalted on MCX, concentrated by vacuum centrifugation and reconstituted in 40 μL 0.1% (v/v) formic acid. The peptide mixture of each sample was labeled using TMT reagent and mixed equally. The labeled peptides were then fractionated by High pH Reversed-Phase Peptide Fractionation Kit (Thermo Scientific). The peptide mixture was reconstituted and acidified with 0.1% TFA solution and loaded to the equilibrated, high-pH, reversed-phase fractionation spin column. Peptides were bound to the hydrophobic resin under aqueous conditions and desalted by washing the column with water by low-speed centrifugation. A step gradient of increasing acetonitrile concentrations in a volatile high-pH elution solution was then applied to the columns to elute bound peptides into 10/20/30 different fractions collected by centrifugation. Finally, the collected fractions of peptides were vacuum dried and lyophilized with 12 μL 0.1% FA.

LC-MS/MS analysis was performed on a Q Exactive mass spectrometer (Thermo Scientific) coupled to Easy nLC (Thermo Scientific). The columns were a reverse phase trap column (Thermo Scientific Acclaim PepMap100, 100 μm*2 cm, nanoViper C18) and a C18- reversed phase analytical column (Thermo Scientific Easy Column, 10 cm long, 75 μm inner diameter, 3 μm resin). The flow rate of the mobile phase (A=0.1% formic acid/water; B=84% acetonitrile and 0.1% formic acid) was 300 nL/min. The mass spectrometer was operated in positive ion mode. MS data was acquired using a data-dependent top20 method that dynamically chose the most abundant precursor ions from the survey scan (300–1800 m/z) for HCD fragmentation. Automatic gain control (AGC) target was set to 1e6, and maximum injection time to 50 ms. Dynamic exclusion duration was 30.0 s. Survey scans were acquired at a resolution of 60,000 at m/z 200 and resolution for HCD spectra was set to 15,000 at m/z 200, and isolation width was 1.5 m/z. Normalized collision energy was 30 eV and the underfill ratio was set as 0.1%. The MS instrument was run with peptide recognition mode enabled.

The MS raw data was searched using MASCOT (Matrix Science, London, UK; version 2.2) ^58^ for identification and quantitation of proteins. MASCOT matched tandem mass spectra to a protein sequence database of *Lactiplantibacillus plantarum* WCFS1 obtained from the Uniprot database ^59^. The protein mass ratio was calculated as the median of only unique peptides of the protein. The quantitative proteomic data can be found in https://github.com/SizheQiu/LbPtEPS/blob/main/data/Proteomics/Proteomics_B.xlsx. Next, gene ontology (GO) and Kyoto Encyclopedia of Genes and Genomes (KEGG) ^51^ annotations were performed for identified proteins using Blast2GO ^60^ and eggNOG-mapper ^61^. The protein subcellular localization was predicted using CELLO ^62^. Differential expression analysis was performed on proteomics data at different pH values with PyDESeq2 ^63^.

### 4.6 Regulatory proteome constrained flux balance analysis

The GSMM used in this study was the modified iBT721 ^20^, and FBA ^19^ was used to compute metabolic fluxes (**see SI, Supplementary method 1.1**). The objective function to maximize was the growth rate normalized to 1 gram dry weight (gDW) of biomass, *v*_growth_ (Eq. 1). The basic constraints were mass conservation (Eq. 2) and default lower/upper bounds (*v*_lb_, *v*_ub_) of reaction fluxes (Eq. 3). COBRApy ^64^ was used to perform FBA.

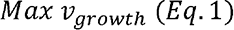

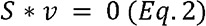

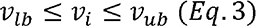

Proteome constrained FBA tightens the metabolic flux solution space by integrating proteome constraints of reactions into conventional FBA ^65^. In this work, the reaction flux (*v_i,_* 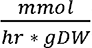) was constrained by the enzyme activity, 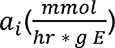 (Eq. 4). Some values of *a_i_* (e.g., Ribose-5-phosphate isomerase) were missing in BRENDA and SABIO-RK, and thus were estimated by DLTKcat ^66^ (**see SI, Supplementary method 1.2**). The proteome of LP HMX2 was divided into sectors of inflexible housekeeping (Q), anabolism (A), transportation (T), catabolism (C) and secondary metabolism (U, EPS biosynthesis). The upper bound of the summation of the 4 flexible sector fractions (*ϕ*_c_ + *ϕ*_A_ + *ϕ*_T_ + *ϕ*_U_) was assumed to be 50% of the total proteome (Eq. 5 and 6) ^43,67,68^. Φ represented the mass fraction of the sector 1 for 1 = A, c, T, u. pTOT was the total mass of the proteome normalized to 1 gDW of biomass 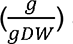 and set as 0.299 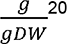 for LP.

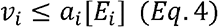

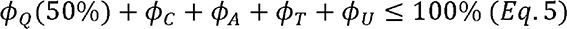

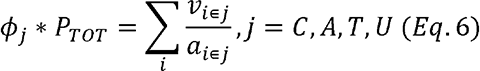

To account for the acid stress induced proteome trade-off between primary and secondary metabolism, the ratio between *ϕ*_U_ and *ϕ*_c_ + *ϕ*_A_ was modeled as a sigmoid function of the pH value (Eq. 7) ^22^. *R* was the base ratio at pH 6.5, and *r*, *k* and *k* were derived based on the quantitative proteomic analysis (**section 4.5**). In addition, the changes in enzyme activities caused by the decrease of pH were considered with the pH-dependent function *F*_pH_ (Eq. 8 & 9). The *F*_pH_ of A, C, T sectors were approximated using the sigmoid function, while that of the U sector was approximated using the quadratic function (**see SI, Supplementary method 1.2**).

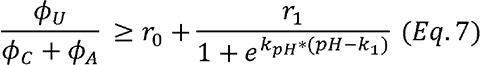

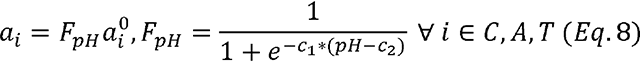

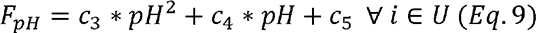

## Acknowledgements

The authors would like to acknowledge the financial support provided by National Natural Science Foundation of China (project No.32302265) and the National Center of Technology Innovation for Dairy (project No. 2023-QNRC-2 and project No. 2023-JSGG-21).

## Author contributions

Sizhe Qiu conceptualized the study, developed the methodology, conducted experiments, curated data, performed data analysis, and contributed to the writing of the manuscript. Aidong Yang contributed to methodology development, writing, and manuscript review. Xinyu Yang contributed to methodology development, experimental work, and writing. Wenlu Li participated in the writing process and manuscript review. Hong Zeng provided input on writing, reviewed the manuscript, and supervised the project. Yanbo Wang contributed to writing, reviewed the manuscript, and provided project supervision.

## Data availability statement

The code and data are openly available at https://github.com/SizheQiu/LbPtEPS.

## Abbreviations

ABC: ATP-binding cassette
BGC: biosynthetic gene cluster
DW: dry weight
EPS: exopolysaccharide
FBA: flux balance analysis
GC: gas chromatography
GSMM: genome scale metabolic model
GT: glycosyltransferase iModulon: a set of independently modulated genes
LC: liquid chromatography
LFC: log2 fold change
LP: *Lactiplantibacillus plantarum*
MRS: De Man, Rogosa and Sharpe
MS: mass spectrometry
PCA: principal component analysis
RPCFBA: regulatory proteome constrained flux balance analysis
RSM: response surface modeling
TF: transcriptional factor

