## Supplementary material for "Proteome trade-off between primary and secondary metabolism shapes acid stress induced bacterial exopolysaccharide production": Table S1, Table S2, Table S3, Figure S1, Figure S2, Figure S3, Figure S4

### Authorship

Sizhe Qiu^1,2^, Aidong Yang^2^, Xinyu Yang^1^, Wenlu Li^1^, Hong Zeng^1,3*^, Yanbo Wang^1*^

^1^School of Food and Health, Beijing Technology and Business University, 100048, China

^2^Department of Engineering Science, University of Oxford, OX1 3PJ, United Kingdom

^3^National Center of Technology Innovation for Dairy, China

### 1. Supplementary method

#### 1.1 Genome-scale metabolic model modification

The GSMM of LP HMX2 was built via the modification of iBT721^1^. This study assumed that the glucose uptake of LP was carried out by D-glucose transport via PEP:Pyr PTS (GLCpts), and hence, GLCt2r was deleted from the original GSMM. Phosphoketolase (PKL), Fructose 6-phosphate aldolase (F6PA) and Dihydroxyacetone phosphotransferase (DHAPT) were removed due to redundancy. The directions of irreversible reactions were all fixed. For reversible reactions (i.e., RPE, RPI, PGI, UDPG4E and MAN6PI), reverse and forward reactions were created. Because GapMind ^2^ found that the amino acid auxotrophy of LP WCFS1 and LP HMX2 were the same, amino acid metabolism was not altered in this GSMM.

Mannose-1-phosphate guanylyltransferase (MAN1PT) was added to the GSMM for the biosynthesis of GDP-Mannose.The EPS biosynthetic pathway was separated as 3 pseudo reactions (i.e., MANT_EPS, GLCT_EPS, and GALT_EPS) and WZX flippase, as shown below:

$MANT\_EPS: 14 GDPMannose[c]\to Man14[c]+ 14 GDP[c]$

$GLCT\_EPS: 6 UDPGlucose[c] + Man14[c] \to Man14GLC6[c]+ 14 UDP[c]$

$GALT\_EPS: UDPGalactose[c] +Man14GLC6[c]\to EPS[c]+UDP[c]$

$WZX: EPS[c]+ ATP[c]+H2O[c]\to EPS[e]+ADP[c]+Pi[c]+H[c]$

$Man14[c]$ and $Man14GLC6[c]$ were pseudo intermediates of the biosynthesis of the pseudo EPS repeating unit, $EPS[c]$. The chemical formulas of $Man14[c]$, $Man14GLC6[c]$ and $EPS[c]$ were C84H154O70, C120H220O100 and C126H231O105, respectively. $[c]$ represented the cytoplasm, and $[e]$ represented the extracellular space.

#### 1.2 Estimation of essential parameters for proteome constrained FBA

For missing enzyme activities ($a_{i}$), DLTKcat^3^ was used to predict $k_{cat}$ for the reaction first, and then the $k_{cat}$(1/s) was converted to $a_{i}$ (mmol/hr*g E) using Eq. S1. The input variables of DLTKcat were substrate SMILES strings, protein sequences and the constant growth temperature, 37 ℃. The detailed user guide of DLTKcat can be found at <https://github.com/SizheQiu/DLTKcat>.

$v_{max}=a_{i}[E_{i}]=k_{cat}\frac{[E_{i}]}{MW_{i}} \Rightarrow a_{i}=\frac{k_{cat}}{MW_{i}} (Eq. S1)$

The activities of PMANM, RPI (reverse) and RPE (reverse) were predicted in this study. For RPI (reverse) and RPE (reverse), the substrates were , respectively.

The mathematical functions of $F_{pH}$ were estimated based on approximated relative activities of enzymes in primary metabolism and EPS biosynthesis (**Figure S2**). In Eq. 8, $c_{1}=1.3812, c_{2}=4.3315$. In Eq. 9, $c_{3}=-0.3815, c_{4}=4.2847, c_{5}=-11.0359$. The average relative enzyme activities for primary metabolism were approximated using normalized growth rates at different pH values. To approximate the relative enzyme activities for EPS biosynthesis, measured EPS production fluxes were divided by total expressions of EPS biosynthetic proteins, and the results were normalized. The enzyme activities of glycosyltransferases (MANT_EPS, GLCT_EPS and GALT_EPS) were estimated based on the maximum EPS production flux at pH 5.5 without the inhibition of undissociated lactic acid (**Figure S3**).

In the simulation, the upper bounds of amino acid uptake fluxes were all set as 0.2 $\frac{mmol}{gDW*hr}$, approximated based on the concentrations of 19 essential amino acids measured during the fermentation (**Figure S4**). As the MRS medium is a rich medium, this study did not set constraints on the uptake fluxes of purines, pyrimidines, metal ions, and vitamins.

### 2. Tables

| **Table S1. Molecular structural properties of LP-HMX2-EPS** | | | |
| --- | --- | --- | --- |
| Monosaccharide | Mass fraction (%) | Glycosyl linkage | Molar fraction (%) |
| Man | 44.69666 | t-Manp | 36.05515 |
| Glc | 40.90727 | 1,2,6-Manp | 15.47439 |
| Gal | 8.579008 | 1,2-Manp | 14.88804 |
| GlcN | 5.602263 | t-Glcp | 13.86694 |
| GalA | 0.17621 | 1,4-Galp | 4.621861 |
| Ara | 0.038584 | 1,4-Glcp | 4.404261 |
| _ | _ | 1,6-Glcp | 3.758698 |
| _ | _ | 1,3-Glcp | 3.676406 |
| _ | _ | t-Galp | 1.150325 |
| _ | _ | t-Fucp | 0.560417 |
| _ | _ | 1,3,4-GalAp | 0.452948 |
| _ | _ | 1,6-Galp | 0.404385 |
| _ | _ | 1,3,6-Glcp | 0.365454 |
| _ | _ | 1,3,6-Manp | 0.320716 |

| **Table S2. Metabolic enzyme information** | | |
| --- | --- | --- |
| ID | Name | EC number |
| GLCpts | D-glucose transport via PEP:Pyr PTS | _ |
| MANpts | D-mannose transport via PEP:Pyr PTS | _ |
| LCTSt | Lactose transport via proton symport | _ |
| LACZ | Beta-galactosidase | 3.2.1.23 |
| GALK | Galactokinase | 2.7.1.6 |
| GALT | Galactose 1 phosphate uridylyltransferase | 2.7.7.10 |
| HEX | Hexokinase (D-glucose:ATP) | 2.7.1.2 |
| PGI | Glucose-6-phosphate isomerase | 5.3.1.9 |
| PFK | Phosphofructokinase | 2.7.1.11 |
| FDPA | Fructose-bisphosphate aldolase | 4.1.2.13 |
| TPI | Triose-phosphate isomerase | 5.3.1.1 |
| GAPD | Glyceraldehyde-3-phosphate dehydrogenase | 1.2.1.12 |
| PGK | Phosphoglycerate kinase | 2.7.2.3 |
| PGM | Phosphoglycerate mutase | 5.4.2.11 |
| ENO | Enolase | 4.2.1.11 |
| PYK | Pyruvate kinase | 2.7.1.40 |
| LDH | Lactate dehydrogenase | 1.1.1.27 |
| PFL | Pyruvate formate lyase | 2.3.1.54 |
| PDH | Pyruvate dehydrogenase | 1.2.7.1 |
| PTAr | Phosphotransacetylase | 2.3.1.8 |
| ACKr | Acetate kinase | 2.7.2.1 |
| ACCOAC | Acetyl-CoA carboxylase | 6.4.1.2 |
| MAN6PI | Mannose-6-phosphate isomerase | 5.3.1.8 |
| PMANM | Phosphomannomutase | 5.4.2.8 |
| MAN1PT | Mannose 1 phosphate guanylyltransferase | 2.7.7.13 |
| PGMT | Phosphoglucomutase | 5.4.2.2 |
| GALU | UTP-glucose-1-phosphate uridylyltransferase | 2.7.7.9 |
| UDPG4E | UDP-glucose 4-epimerase | 5.1.3.2 |
| G6PDH | Glucose 6-phosphate dehydrogenase | 1.1.1.49 |
| PGL | 6-phosphogluconolactonase | 3.1.1.31 |
| GND | Phosphogluconate dehydrogenase | 1.1.1.44 |
| RPE | Ribulose 5-phosphate 3-epimerase | 5.1.3.1 |
| RPI | Ribose-5-phosphate isomerase | 5.3.1.6 |
| TKT1/2 | Transketolase | 2.2.1.1 |
| TALA | Transaldolase | 2.2.1.2 |

* FBA (fructose-bisphosphate aldolase) was renamed as FDPA in this study to avoid confusion with FBA (flux balance analysis).

* Detailed enzyme and reaction information can be found in the BIGG database^4^.

| **Table S3. Enzyme specific activity values** | | |
| --- | --- | --- |
| Enzyme | $a_{i}$ ($\frac{mmol}{hr * g E}$) | Source |
| LCTSt | 540 | ^5^ |
| GLCpts/MANpts | 361.14 | ^6^ |
| HEX | 9480 | ^7^ |
| G6PDH2r | 6240 | ^8^ |
| PGL | 31200 | ^9^ |
| GND | 1920 | ^10^ |
| RPE | 133886.21 (+)/133.94 (-) | ^11^/Predicted by DLTKcat |
| RPI | 316635.95 (+)/4793.05 (-) | ^12^/Predicted by DLTKcat |
| PGI | 12756 (+)/870 (-) | ^13^/^14^ |
| PFK | 11400 | ^15^ |
| FDPA | 28620 | ^16^ |
| GAPD | 2400 | ^17^ |
| PGK | 28800 | ^18^ |
| PGM | 7440 | ^19^ |
| ENO | 15600 | ^20^ |
| PYK | 3300 | ^21^ |
| LDH_L | 141000 | ^22^ |
| LDH_D | 126000 | ^23^ |
| ACALD | 2916 | ^24^ |
| ALCD2x | 11451.6 | ^25^ |
| PFL | 720 | ^26^ |
| PDH | 1500 | ^27^ |
| PTAr | 428400 | ^28^ |
| ACKr | 65220 | ^29^ |
| ACCOAC | 360 | ^30^ |
| PGMT | 1890 | ^31^ |
| UDPG4E | 1620 (+)/13998 (-) | ^32^/^33^ |
| GALU | 285.6 | ^34^ |
| GALK | 690 | ^35^ |
| GALT | 1020 | ^36^ |
| MAN6PI | 270 (+)/ 639.79 (-) | ^37^/^38^ |
| PMANM | 82.99 | Predicted by DLTKcat |
| MAN1PT | 144 | ^39^ |
| MANT_EPS/GLCT_EPS/GALT_EPS | 250.61 | Estimated in this study |
| WZX | 37.44 | ^40^ |
| Growth (biomass_LPL60) | 107.4 | ^41^ |
| Acid exportation (lactic and acetic acid) | 6360 | ^42^ |

* (+) means the forward reaction and (-) means the reverse reaction.

### 3. Figures

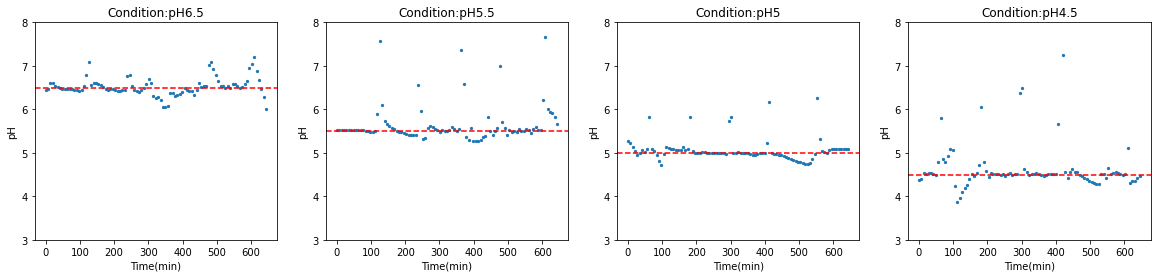

Figure S1. The control of pH for 4 different conditions: pH=6.5, 5.5, 5.0 and 4.5.

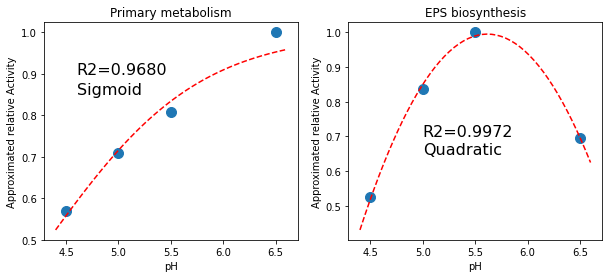

Figure S2. The approximation of pH dependent enzyme activity coefficients for enzymes in primary metabolism (left) and EPS biosynthesis (right).

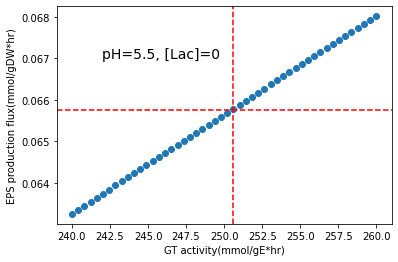

Figure S3. The estimation of enzyme activities of glycosyltransferases (MANT_EPS, GLCT_EPS and GALT_EPS).

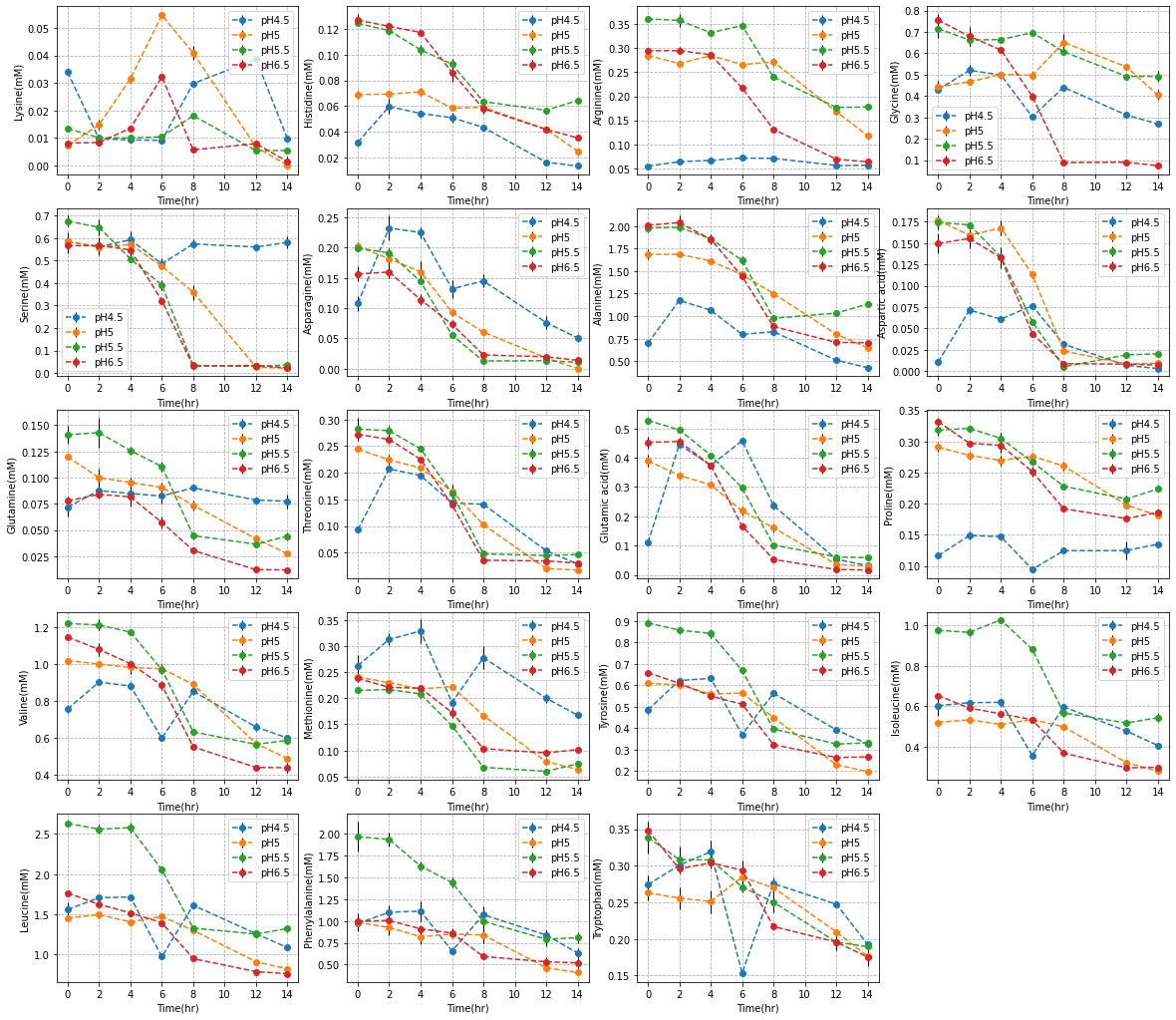

Figure S4. The concentrations of 19 essential amino acids. Cysteine was not detected.
